# Whole Genome Sequencing for Revealing the Point Mutations of SARS-CoV-2 Genome in Bangladeshi Isolates and their Structural Effects on Viral Proteins

**DOI:** 10.1101/2020.12.05.413377

**Authors:** Mohammad Uzzal Hossain, Ishtiaque Ahammad, Md. Tabassum Hossain Emon, Arittra Bhattacharjee, Zeshan Mahmud Chowdhury, Md. Golam Mosaib, Keshob Chandra Das, Chaman Ara Keya, Md. Salimullah

**Author notes:** **Correspondence:** Dr. Md. Salimullah, Chief Scientific Officer, Molecular Biotechnology Division, National Institute of Biotechnology, Ganakbari, Ashulia, Savar, Dhaka-1349, Bangladesh.

## Abstract

Coronavirus disease-19 (COVID-19) is the recent global pandemic caused by the virus Severe Acute Respiratory Syndrome Coronavirus-2 (SARS-CoV-2). The virus has already killed more than one million people worldwide and billions are at risk of getting infected. As of now, there is neither any drug nor any vaccine in sight with conclusive scientific evidence that it can cure or provide protection against the illness. Since novel coronavirus is a new virus, mining its genome sequence is of crucial importance for drug/vaccine(s) development. Whole genome sequencing is a helpful tool in identifying genetic changes that occur in a virus when it spreads through the population. In this study, we performed complete genome sequencing of SARS-CoV-2 to unveil the genomic variation and indel, if present. We discovered thirteen (13) mutations in Orf1ab, S and N gene where seven (7) of them turned out to be novel mutations from our sequenced isolate. Besides, we found one (1) insertion and seven (7) deletions from the indel analysis among the 323 Bangladeshi isolates. However, the indel did not show any effect on proteins. Our energy minimization analysis showed both stabilizing and destabilizing impact on viral proteins depending on the mutation. Interestingly, all the variants were located in the binding site of the proteins. Furthermore, drug binding analysis revealed marked difference in interacting residues in mutants when compared to the wild type. Our analysis also suggested that eleven (11) mutations could exert damaging effects on their corresponding protein structures. The analysis of SARS-CoV-2 genetic variation and their impacts presented in this study might be helpful in gaining a better understanding of the pathogenesis of this deadly virus.

## Introduction

COVID-19 can be currently considered an menace to humankind brought about by the novel Severe Acute Respiratory Syndrome Coronavirus-2 (SARS-CoV-2) which began its journey from the Wuhan province of the People’s Republic of China. [1]–[8] The infection basically targets the respiratory framework of its host causing influenza-like sickness with symptoms such as cough, fever, and in progressively serious cases, troubled breathing. [9]–[15] According to the data available, mortality is higher in individuals of advanced ages (> 60 years) and the ones with comorbidities [16]–[21]. Apart from intense respiratory problems, COVID-19 has been shown to cause systemic irritation prompting sepsis [22]–[25], intense cardiovascular injury [26]–[31], cardiovascular breakdown [26], [32]–[36] and multiorgan failure in critical patients. [37] COVID-19 has been rightly announced as a global pandemic by the World Health Organization (WHO) as it has spanned over 200 countries and territories around the world. As of Oct 20, 2020, more than 40,648,527 individuals have been infected up until this point, with around 9,172,183 active cases and 31,476,344 closed cases and 1,122,992 deaths (https://www.worldometers.info/coronavirus/). While the USA has the highest number of confirmed cases (with over 8,456,653 so far) and has more than 225,222 deaths to date. [38] China, the nation where everything started in November, is presently on the downslope sustaining just above 200 active cases. [39]–[41] The vast majority of the countries around the world were in lockdown to stop human to human transmission and halt the spread of the disease. [42], [43]

Coronaviruses (CoVs) are enveloped, single-stranded, positive RNA viruses that are pathogenic to their hosts. [44]–[48] SARS-CoV-2 is the causative agent behind COVID-19 and is more pathogenic in contrast with previously observed SARS-CoV (2002) and Middle East respiratory syndrome coronavirus (MERS-CoV, 2013)[49]–[58]. There is a dire need to examine the virus more comprehensively to comprehend the system of pathogenesis, its destructiveness to develop powerful therapeutic measures. [59] CoVs belongs to the Coronaviridae family under Nidovirales order. They have been grouped into four genera that belong to α-, β-, γ-, and δ-coronaviruses. [60] Among them, α-and β-COVs infect vertebrates, γ-coronaviruses avians, while the δ-coronaviruses infect both. SARS-CoV, mouse hepatitis coronavirus (MHV), MERS-CoV, Bovine coronavirus (BCoV), bat coronavirus HKU4, and human coronavirus OC43, including SARS-CoV-2, are β-coronaviruses. [61] Zoonotic transmission is the medium of transmission for each of the three CoVs, SARS-, MERS-, and SARS-CoV-2, and they spread through close contact. The essential multiplication number (R0) of the individual-to-individual spread of SARS-CoV-2 is around 2.6, which implies that the confirmed cases develop at a striking exponential rate. [62] CoVs being 26 to 32 kb long have the biggest RNA viral genome. [63] The SARS-CoV-2 genome share approximately 90% identity with essential enzymes and structural proteins of SARS-CoV. Fundamentally, SARS-CoV-2 contains four basic proteins known as-spike (S), envelope (E), membrane (M), and nucleocapsid (N) proteins. These proteins share high sequence similarity with the sequence of the corresponding proteins in SARS– CoV, and MERS-CoV. Hence, it is vital to scrutinize the SARS-CoV-2 genome to determine why this infection is progressively inclined to be more infectious and lethal than its predecessors.

Utilizing Sanger sequencing and cutting-edge whole genome sequencing of SARS-CoV-2 isolates from oropharyngeal samples, we depicted the genomic portraits of two genomes alongside other Bangladeshi strains. [64]

In this study, we have analyzed the genomic arrangements of SARS-CoV-2 to identify the mutations found within the genomes and anticipate their effect on the protein structure from a structural biology perspective in order to shed light on the suitable therapeutics against this deadly virus.

## Methods

### Virus Isolation

The patient’s oropharyngeal samples SARS-CoV-2/human/BGD/NIB_01/2020 and SARS-CoV-2/human/BGD/NIB-BCSIR_02/2020 were collected using the UTM™ kit containing 1 mL of viral transport media (Copan Diagnostics Inc., Murrieta, CA, USA) on day 7 of the patient’s illness with symptoms of cough, mild fever, and throat congestion. The specimen was tested positive for SARS-CoV-2 by real-time reverse transcriptase PCR (rRT-PCR). Then, the viral RNA was extracted directly from the patient’s swab using PureLink Viral RNA/DNA Mini Kit (Invitrogen). The viral RNA was then converted into cDNA using SuperScript™ VILO™ cDNA Synthesis Kit (Invitrogen) according to the manufacturer’s instructions.

### DNA Sequencing

#### Sanger Dideoxy sequencing

The forty eight (48) pair primers were designed to cover the whole genome of the virus by following two conditions: (1) their sequence is conserved among all the available SARS-CoV-2 isolates and (2) the terminal of the amplicons will overlap with neighboring amplicons. The polymerase chain reaction (PCR) was performed and the 48 primers then generated 47 amplicons which were visualized in 1.5% Agarose gel electrophoresis. The amplicons were further purified using Purelink PCR purification kit (ThermoFisher Scientific, USA). These purified amplicons were sequenced using Sanger dideoxy method by “ABI 3500” with BigDye Terminator version 3.1 cycle sequencing kit (Applied Biosystems, USA). The raw reads were assembled by DNA Baser (https://www.dnabaser.com) [65] and verified by SeqMan Pro®. Version 14.1. DNASTAR. [66] Madison, WI. These overlapping regions were visualized by CLC Genomics Workbench 20.0.4 (https://digitalinsights.qiagen.com) and merged with EMBOSS: merger (https://www.bioinformatics.nl/cgi-bin/emboss/merger).

#### NGS Sequencing

Illumina Nextseq 550 next-generation sequencing technology was used to sequence the complete genome of the SARS-CoV-2/human/BGD/NIB-BCSIR_02/2020 isolate to where Nextera DNA Flex was utilized as library preparation kit for the synthesis of the nucleotides. [67] To cover the 300 cycle, the NextSeq High Output Kit was utilized as the reagent cartridge. To generate the FASTQ data workflow the run mode was set as local run manager in every NextSeq 4-channel chemistry. Analysis and Quality check was performed using a customized version of the DRAGEN RNA pipeline, which was also available on local DRAGEN server hardware. The Illumina® DRAGEN RNA Pathogen Detection App uses a combined human and virus reference to analyze pathogen data. The raw reads were cleaned by trimming low-quality bases with Trimmomatic 0.36 (-phred33, LEADING:20, TRAILING:20, SLIDCitation. The assembly was performed by the utilization of SPAdes using default parameters as well as used to cross-validate with the reference-based method as an internal control.

#### Variant Identification

Basic Local Alignment Search Tools (BLAST) [68] was employed to identify possible mutations in Sanger Based sequenced nucleotide sequences. Nucleotide program of blast was selected for this identification. The mapped polymorphisms were investigated for their occurrence frequency worldwide and checked for their profile at CNCB^3^ resource. Chimera was utilized to visualize the mapped polymorphisms. Besides, all the available Bangladeshi strains (n=323) of SARS-CoV-2 were retrieved from GISAID [69] and further explored to find out the most common mutations.

#### Indel Analysis

We ran BLAST for all Bangladeshi SARS CoV-2 genome profiles, to get a view on the pairwise alignment comparison. Deleted sequences were closely investigated in Artemis Comparison Tool window. [70] Gene regions are observed discreetly.

#### Mutational Effect Analysis

To observe the mutational effect of mapped polymorphisms, 3 dimensional (3D) structure was built through the ROBETTA prediction server. [71] Later, the difference of the energy was calculated by GROMACS in both wild type and mutant 3D structures to estimate the structural abnormality and change in stability. [72] Binding site of both wild and mutant structure was analyzed to check whether the amino acid residues are into the binding site region or not.

#### Drug-Binding Analysis

We have performed Molecular Docking using Autodock vina[73] for the analysis of interacting residues to the druggable targets. We retrieved the structures of all the interacting drugs (.pdb files) by virtual screening of the Drugbank database. Then we generated the .pdbqt files of the targets of polymorphisms for docking experiments. Blind docking was performed for the identification of the most effective binding site of these drugs. A grid box parameter for covering the whole protein was set for all docking runs.

## Results

### Revealing the SARS-CoV-2 Genome

The workflow of this manuscript has been shown in **Fig 1.**

**Fig 1:**
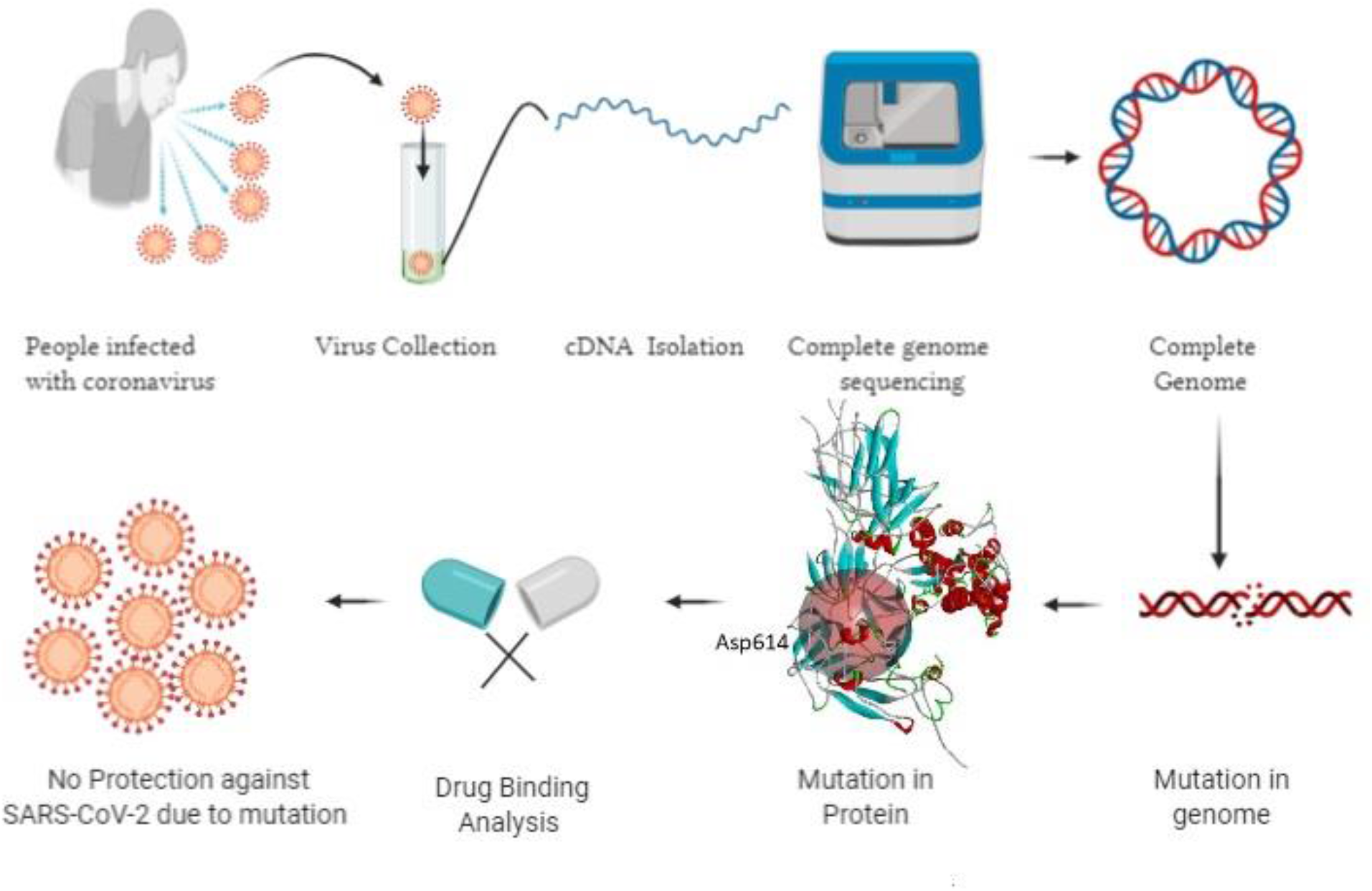
Complete workflow of the study.

In case of NIB-01 isolates forty eight (48) contigs with ninety four (94) overlapping regions were obtained. The sequence coverage was 2X the raw reads. It was then assembled by SeqMan Pro and EMBOSS merger. [74] The assembled viral genome consisted of a single stranded positive (+) RNA that is 29,724 nucleotides long with: 8,882 Adenosines (29.88%), 5,455 Cytosine (18.35%), 5,836 Guanine (19.63%), and 9,551 Thymine (32.13%). The GC content of the whole genome was 38%. A total of 1,78,22,898 reads were produced in the reference-based alignment after trimming 99% of them were mapped to the SARS-CoV-2 reference genome.

The complete nucleotide sequence of SARS-CoV-2 isolate SARS-CoV-2/human/BGD/NIB_01/2020 from the Sanger sequencing has been deposited in GenBank under the accession number MT509958 (https://www.ncbi.nlm.nih.gov/nuccore/MT509958).The complete nucleotide sequence of SARS-CoV-2/human/BGD/NIB-BCSIR_02/2020 isolate from the NGS has been also deposited under the accession number MT568643 (https://www.ncbi.nlm.nih.gov/nuccore/MT568643.1?report=genbank).

### Exploration of variant in the sequenced isolates

We have found thirteen (13) mutations at the 93^rd^; C → T, 479^th^; T → A, 481^th^; C→A; 1015^th^; A → T, 2889^th^; C → T, 5098^th^; G → T; 5237^th^; C → T, 5642^th^; G → T, 8023^rd^; G → A, 23255^th^; A → G 28733^rd^; G → A, 28734^th^; G → A and 28735^th^; G → C from the MT509958 whole genome (**Table 1**). However, the MT568643 whole genome showed no mutation against the reference sequence. From them, six (6) mutations namely 93^rd^; C → T, 2889^th^; C → T, 23255^th^; A→G, 28733^rd^; G → A, 28734^th^; G → A and 28735^th^; G → C were found in CNCB^3^ resource where the available mutations of SARS-CoV-2 were enlisted (**Table 1**). These mutations have already been found in different countries where the SARS-Cov-2 has been sequenced. These mutations were mostly found in the United States of America (USA) and the United Kingdom (UK) (**Table 1**). The position of 93^rd^; C → T mutation is located in 5’UTR upstream region. And the position 2889^th^; C → T mutation has shown no change to its protein sequence. The other 23255^th^; A→G, 28733^rd^; G → A, 28734^th^; G → A and 28735^th^; G → C mutations can alter the amino acid sequence and can have the missense effect on the protein (**Table 1**). Apart from these 6 mutations, seven (7) mutations have shown as unique variants against the reference sequence (**Table 1 and Fig 2**). There is no report previously having found for these mutations. Besides, we analyzed our assemble genome to look for any insertion/deletions but these two genomes contain no deletions/insertions. However, we have identified one (1) insertion and seven (7) deletions of the eight (8) Bangladeshi strain EPI_ISL_466692, EPI_ISL_450343, EPI_ISL_450344, EPI_ISL_468074, EPI_ISL_514614, EPI_ISL_445213, EPI_ISL_445217 and EPI_ISL_450842 (**Table 2 and Fig 3**). We additionally scrutinized these deleted regions but we didn’t find any domain or motif on this region. Apart from our reported complete genomes, we have also identified the most common mutations occurred in Bangladesh from the complete genomes reported in GSAID database from Bangladesh. These genomes showed three mutations in the positions 14408, 23403 and 28878 compared to reference genome (**Table 1**).

**Table 1:** NIB_01 polymorphisms against reference sequence and their mutational effect.

| SL No . | Query (Reference) Position | Position in subject(NIB-01) sequence | Amino acid | Query → Subject | Gene | Mutants Observed | Mutants observed in No. in Viruses | Impact on Proteins | Protein ID of mutants |
| --- | --- | --- | --- | --- | --- | --- | --- | --- | --- |
| 1. | 241<sup>th</sup> | 93 | - | **C** → T | 5` UTR | US,<br>UK | 1504<br>2 | Upstre<br>am<br>variant | QH<br>D43<br>415.<br>1 |
| 2. | 627<sup>th</sup> | 479 | Val121 | **T** → A | Orf1ab | - | - | - | - |
| 3. | 629<sup>th</sup> | 481 | Leu122 | **C** → A | Orf1ab | - | - | - | - |
| 4. | 1163<sup>th</sup> | 1015 | Ile120 | **A** → T | Orf1ab | - | - | Coding<br>Variant | - |
| 5. | 3037<sup>th</sup> | 2889 | Phe924 | **C** → T | Orf1ab | US,<br>UK | 1505<br>9 | Coding<br>Variant | QH<br>D43<br>415.<br>1 |
| 6. | 5246<sup>th</sup> | 5098 | Val843 | **G** → T | Orf1ab | - | - | Coding<br>Variant | QH<br>D43<br>415.<br>1 |
| 7. | 5385<sup>th</sup> | 5237 | Ala889 | **C** → T | Orf1ab | - | - | Coding<br>Variant | QH<br>D43<br>415.<br>1 |
| 8. | 7790<sup>th</sup> | 5642 | Gly1691 | **G** → T | Orf1ab | - | - | Coding<br>Variant | QH<br>D43<br>415.<br>1 |
| 9. | 8171<sup>th</sup> | 8023 | Ala1818 | **G** → A | Orf1ab | - | - | Coding<br>Variant | QH<br>D43<br>415.<br>1 |
| 10. | 23403<sup>th</sup> | 23255 | Asp614 | **A** → G | “S” | US,<br>UK | 1511<br>8 | Coding<br>Variant | QH<br>D43<br>416.<br>1 |
| 11. | 28881<sup>th</sup> | 28733 | Ser202 | **G** → A | “N” | UK | 4551 | Coding Variant | QH D43 423. 2 |
| 12. | 28882<sup>th</sup> | 28734 | Arg203 | **G** → A | “N” | UK | 4542 | Coding Variant | QH D43 423. 2 |
| 13. | 28883<sup>th</sup> | 28735 | Gly204 | **G** → C | “N” | UK | 4537 | Coding Variant | QH D43 423. 2 |

**Table 2:** Indel profile of Bangladeshi whole genome.

| Serial No. | Sequence Identifier | Indel Type | Region |
| --- | --- | --- | --- |
| 1) | hCoV-19/Bangladesh/BCSIR-NILMRC-071/2020\|EPI_ISL_466692\|2020-05-26 | Insertion | ORF8 (27910-27985) |
| 2) | hCoV-19/Bangladesh/BARJ-CVASU-CTG-511/2020\|EPI_ISL_450343\|2020-05-09 | Deletion | ORF8 (27913-28254) |
| 3) | hCoV-19/Bangladesh/BARJ-CVASU-CTG-517/2020\|EPI_ISL_450344\|2020-05-03 | Deletion | ORF8 (27913-28254) |
| 4) | hCoV-19/Bangladesh/CHRF-0006/2020\|EPI_ISL_468074\|2020-05-17 | Deletion | ORF8<br>(27913-28254) |
| 5) | hCoV-19/Bangladesh/NGRI-NSTU-31/2020\|EPI_ISL_514614\|2020-07-21 | Deletion | ORF8<br>(27913-28254) |
| 6) | hCoV-19/Bangladesh/DNAS-CPH-467/2020\|EPI_ISL_445213\|2020-04-28 | Deletion | ORF7<br>(27476-27668) |
| 7) | hCoV-19/Bangladesh/DNAS-CPH-436/2020\|EPI_ISL_445217\|2020-04-28 | Deletion | ORF7<br>(27476-27668) |
| 8) | hCoV-19/Bangladesh/DU-50761/2020\|EPI_ISL_450842\|2020-05-06 | Deletion | ORF7<br>(27476-27668) |

**Fig 2:**
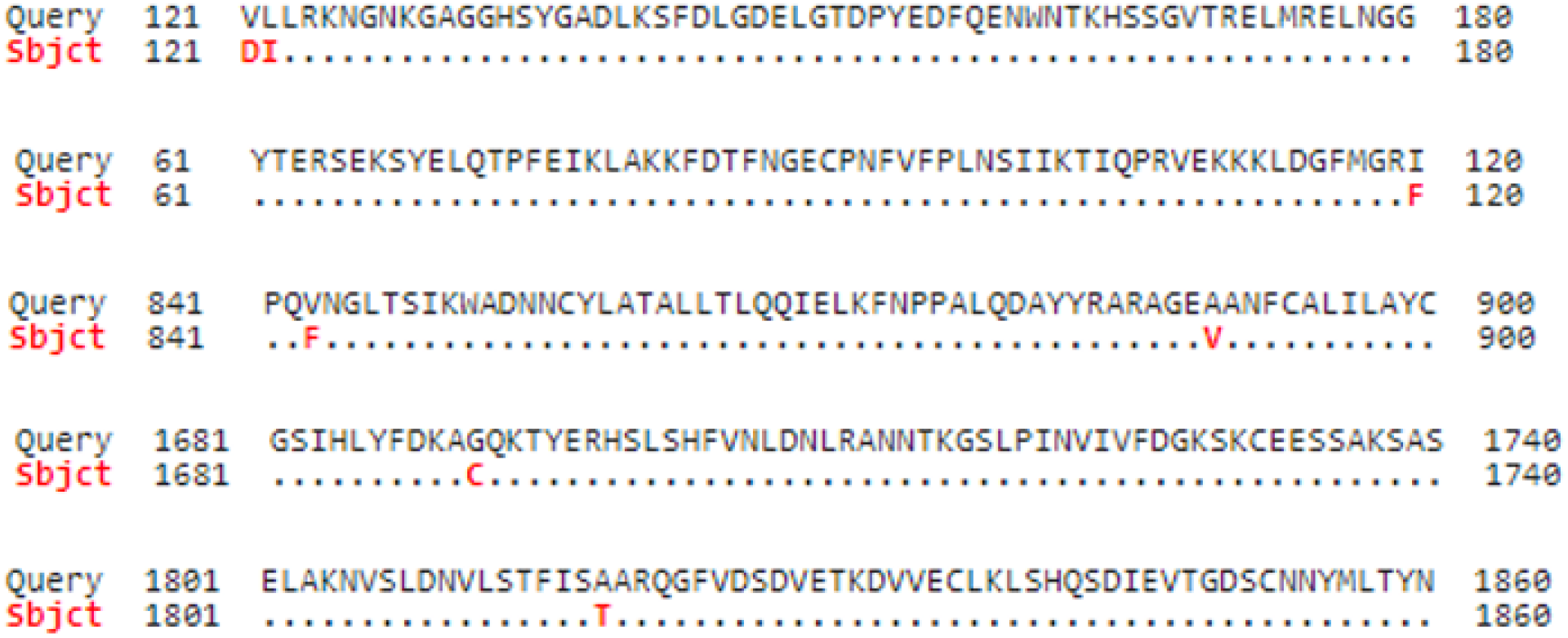
Novel Mutation in protein sequence in NIB_01 whole genome.

**Fig 3:**
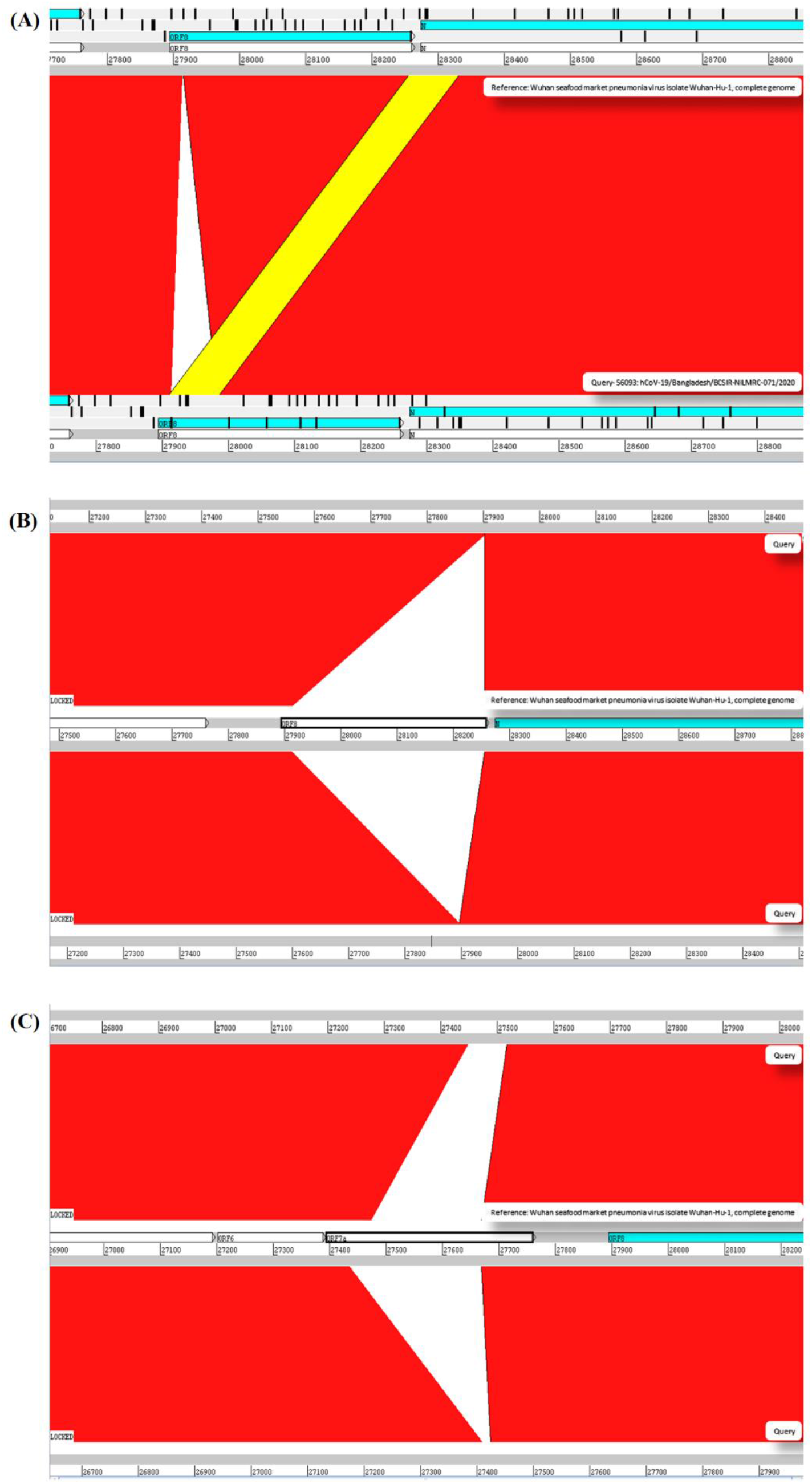
Deletion of sequences in complete genome of the Bangladesh strain. Indel experimentation result from comparative genome browsing against reference genome in Artemis window. A) Mobile element insertion from **N** to **ORF8** in EPI_ISL_466692. Yellow bar representing homologous sequence moiety between these two regions. B) EPI_ISL_450343, EPI_ISL_450344, EPI_ISL_468074, EPI_ISL_514614 are sharing the same deletion pattern in **ORF8**. Majority portion of **ORF8**, lost in the deletion event. C) EPI_ISL_445213, EPI_ISL_445217 and EPI_ISL_450842 are also sharing the same deletion pattern in **ORF7a**. Majority portion of **ORF7a**, lost in the deletion event.

### Point mutations alter respective protein structures

We have analyzed the mutational effect of all the mutations. Therefore, the 3D structure was built to explore the mutational effect on the protein structure (**Figure 4**). In this case, two types of 3D structure was built i) the structure with wild type residue and ii) the structure with mutant residue. We have performed the energy minimization of both the wild type and mutants. We have found significant differences in the stability of the structure upon mutation (**Table 3**). Mutants 479^th^; T → A and 1015^th^; A → T showed higher energy minimization which predicted these proteins to be more stable than the wild type. All the other protein models based on mutation showed less energy minimization than the wild type protein model. Therefore, these protein structures could be more unstable upon mutation in the protein sequence. The highest difference was observed in the mutation in the 5642^th^ position; G → T mutation (from −23276.78 kj/mol to −22377.976 kj/mol) (**Table 3**). Afterwards, the binding site was analyzed to determine whether the wild type and the mutant residues fell within the ligand binding site or not. The binding site residues confirmed that all the mutations from the complete genome belonged to the binding site region (**Fig 5**). Later, we have performed the drug binding analysis followed by virtual drug screening in DrugBank server. Ivermectin and Remdisivir drugs topped the list of potential drug candidates. We then prepared the protein structures and converted them to .pdbqt format for molecular docking experiment. We identified the binding site region for each protein and set the grid box to allow the drugs only to bind to that specific region. The binding affinity analysis showed that compared to the wild type, the drug Ivermectin bound more tightly to proteins which has mutation at positions 479 (T → A), 5642 (G → T) and 8023 (G → A) which bound more loosely to proteins with mutations at positions 481 (C→A), 1015 (A → T), 5098 (G → T) and 5237 (C → T). Remdisivir binds more tightly to proteins with mutations at 28733 (G → A), 28734 (G → A), and 28735 (G → C) and more loosely to 23255 (A → G) compared to the wild type (**Table 4**). It is to be noted that the interaction of residues of wild type protein were found to be different than that of mutant model. For example, 479^th^; T → A mutant model which acquired the V→D amino acid interacted with Glu37, Glu41, LEU177, GLY180, LEU104, VAL108, HIS110, Glu87, Leu88, Lys141, Tyr154 residues whereas wild type interacted with Leu18, Val28, Glu37, Glu41, His 45, Leu53, Val54, Ile71, Arg73, Val86,, Val121, Leu122, Asp139 (**Fig 4**).

**Table 3:**
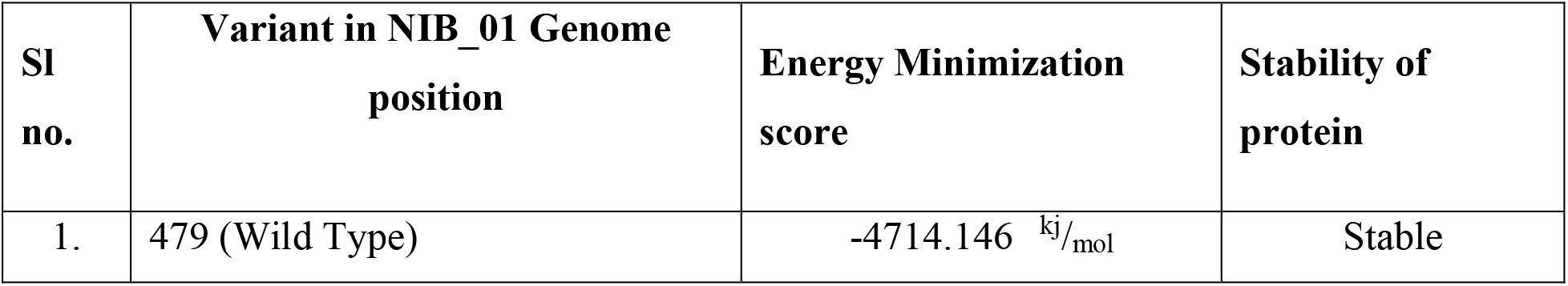
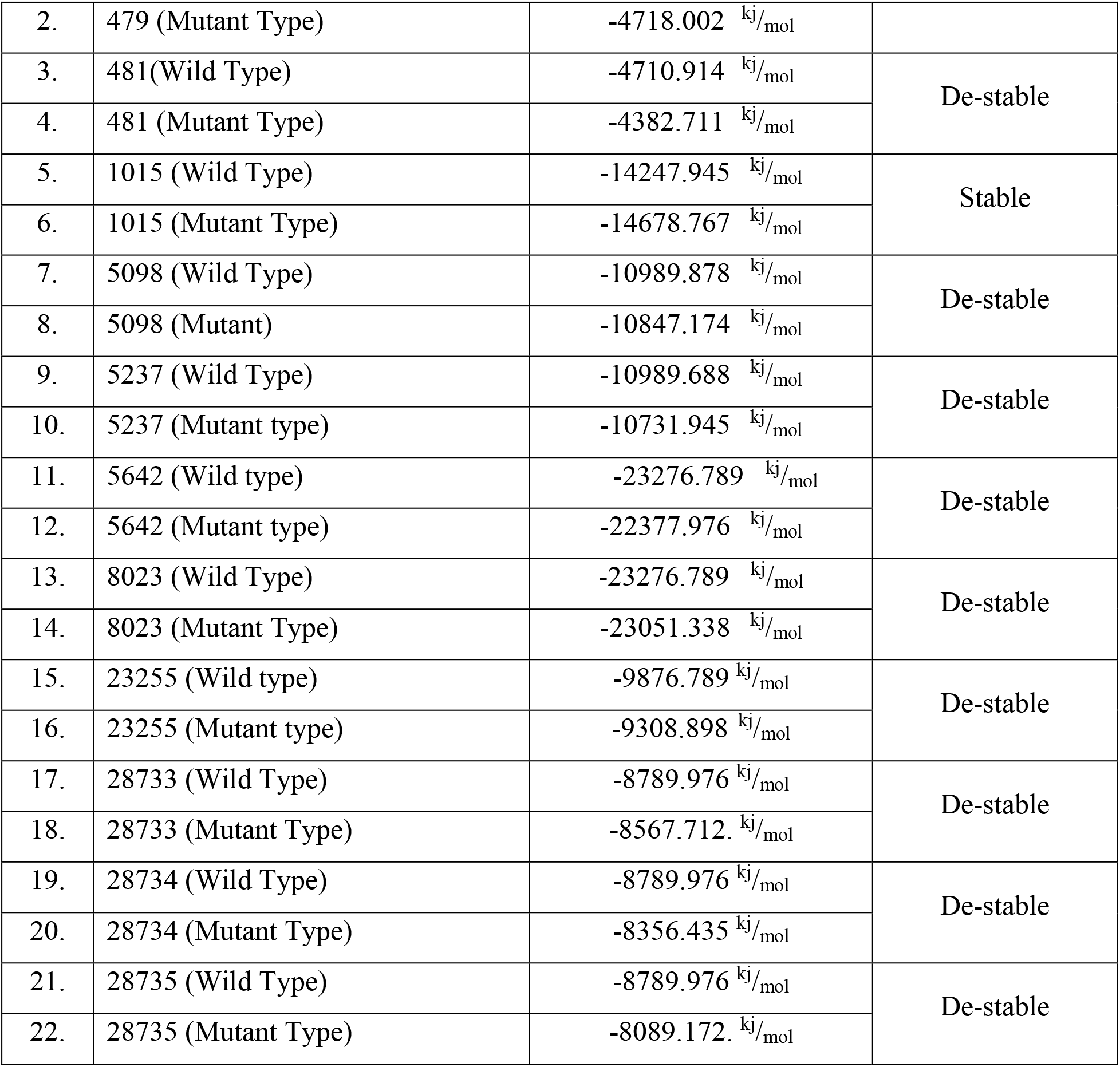
Energy minimization of the wild type and mutant model.

**Table 4:** Drug binding analysis of variant.

| Variant Position<br>in subject(NIB-01) sequence | Drug<br>Name | Binding<br>Energy | Amino Acid Residue |
| --- | --- | --- | --- |
| 479 (Wild Type) | Ivermectin | -7.9 kcal/mol | Leu18, Val28, Glu37, Glu41, His 45,<br>Leu53, Val54, Ile71, Arg73, Val86, ,<br>**Val121**, Leu122, Asp139 |
| 479 (Mutant Type) | Ivermectin | -8.2 kcal/mol | Glu37, Glu41, LEU177, GLY180, LEU104, VAL108, HIS110, Glu87, Leu88, Lys141, Tyr154 |
| 481(Wild Type) | Ivermectin | -8.7 kcal/mol | Leu18, Val28, Glu37, Glu41, His 45, Leu53, Val54, Ile71, Arg73, Val86, , Val121, **Leu122**, Asp139 |
| 481 (Mutant Type) | Ivermectin | -7.7 kcal/mol | Glu37, Glu41, LEU177, GLY180, LEU104, VAL108, HIS110, Glu87, Leu88, Lys141, Tyr154 |
| 1015 (Wild Type) | Ivermectin | -7.9 kcal/mol | Lys22, Met135, His202, Glu206, Lys214, Arg107, **Ile120**, Val126, Asn133, Phe633, Thr634, Ala889 |
| 1015 (Mutant Type) | Ivermectin | -6.9 kcal/mol | GLY235, ALA241, ILE251, GLY260, ASP268, SER303, LYS317, LYS335, GLU359, ALA360, THR371, PHE406, THR412 |
| 5098 (Wild Type) | Ivermectin | -8.8 kcal/mol | ASP137, THR149, TYR170, **VAL173**, CYS186, ALA206, Ala219, CYS230 |
| 5098 (Mutant) | Ivermectin | -8.0 kcal/mol | LEU133, PRO134, SER178, LEU195, ARG213, ALA214, LEU225 |
| 5237 (Wild Type) | Ivermectin | -7.6 kcal/mol | ASP137, THR149, TYR170, VAL173, CYS186, ALA206, **Ala219**, CYS230 |
| 5237 (Mutant type) | Ivermectin | -7.1 kcal/mol | LEU133, PRO134, SER178, LEU195, ARG213, ALA214, LEU225 |
| 5642 (Wild type)) | Ivermectin | -6.7kcal/mol | HIS12, CYS23, VAL33, SER52, LYS61, TYR68, **GLY73**, VAL104, ILE133, SER147, THR196<br>ALA200, ASN235, ARG250 |
| 5642 (Mutant type) | Ivermectin | -8.5 kcal/mol | PHE69, ASN87, ILE105, LYS119, LEU135, VAL150, ALA151, GLU181, ASP251, LEU252, CYS258 |
| 8023 (Wild Type) | Ivermectin | -6.6 kcal/mol | HIS12, CYS23, VAL33, SER52, LYS61, TYR68, GLY73, VAL104, ILE133, SER147, THR196<br>**ALA200**, ASN235, ARG250 |
| 8023 (Mutant Type) | Ivermectin | -7.5 kcal/mol | PHE69, ASN87, ILE105, LYS119, LEU135, VAL150, ALA151, GLU181, ASP251, LEU252, CYS258 |
| 23255 (Wild type) | Remdisivir | -8.7 kcal/mol | Gln321, Glu324, Ile326, His519, Phe543, Lys557, Thr553, Lys 557, Ile569, Ala570, **Asp614** |
| 23255 (Mutant type) | Remdisivir | -8.3 kcal/mol | ASN87, ILE128, ASN234, ILE235, ASP290, PHE329, PHE347, LEU368, ASN394, THR415, GLY416, ASN501, PHE515, GLY526 |
| 28733 (Wild Type) | Remdisivir | -7.7 kcal/mol | Ser21, Asp22, **Ser202**, Arg203, Gly204Leu224, Arg226, Glu280, Gln281, Ser318, Lys338, Val392 |
| 28733 (Mutant Type) | Remdisivir | -8.1 kcal/mol | ASN192, GLY295, GLN306, GLU323, SER327, GLY328, ASP343, LEU353, ASN354, THR366, THR379, VAL392, ASP415, SER416, THR417, GLN418, ALA419 |
| 28734 (Wild Type) | Remdisivir | -6.8 kcal/mol | Ser21, Asp22, Ser202, **Arg203**, Gly204Leu224, Arg226, Glu280, Gln281, Ser318, Lys338, Val392 |
| 28734 (Mutant Type) | Remdisivir | -7.8 kcal/mol | ASN192, GLY295, GLN306, GLU323, SER327, GLY328, ASP343, LEU353, ASN354, THR366, THR379, VAL392, ASP415, SER416, THR417, GLN418, ALA419 |
| 28735 (Wild Type) | Remdisivir | -7.1 kcal/mol | Ser21, Asp22, Ser202, Arg203, Gly204, Leu224, Arg226, Glu280, Gln281, Ser318, Lys338, Val392 |
| 28735 (Mutant Type) | Remdisivir | -8.4 kcal/mol | ASN192, GLY295, GLN306, GLU323, SER327, GLY328, ASP343, LEU353, ASN354, THR366, THR379, VAL392, ASP415, SER416, THR417, GLN418, ALA419 |

Red color denotes the wild mutation.

**Fig 4:**
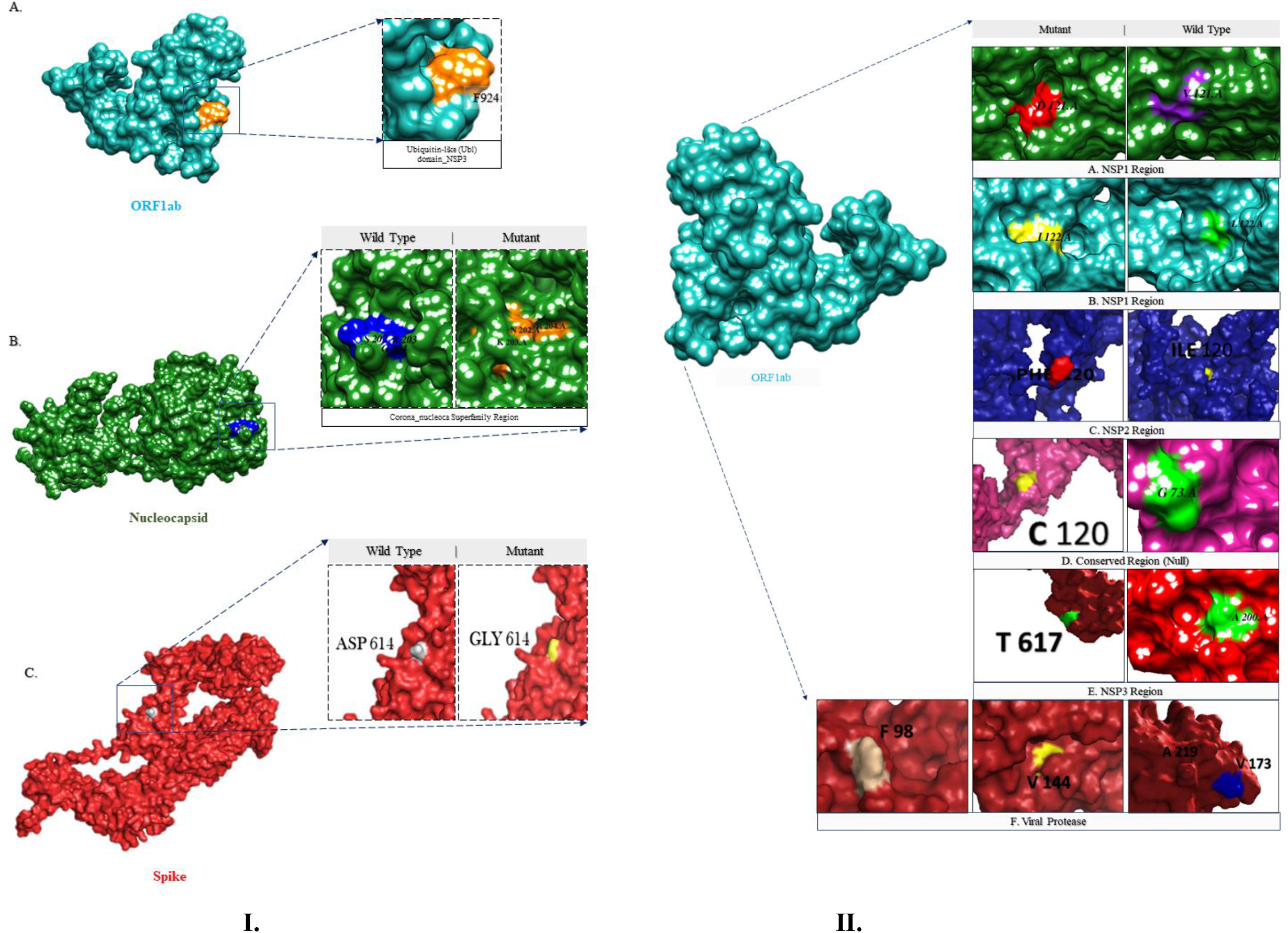
Amino acid location in tertiary structure. I) Previously Found Variants. II) Unique Variants to Bangladesh.

**Fig 5:**
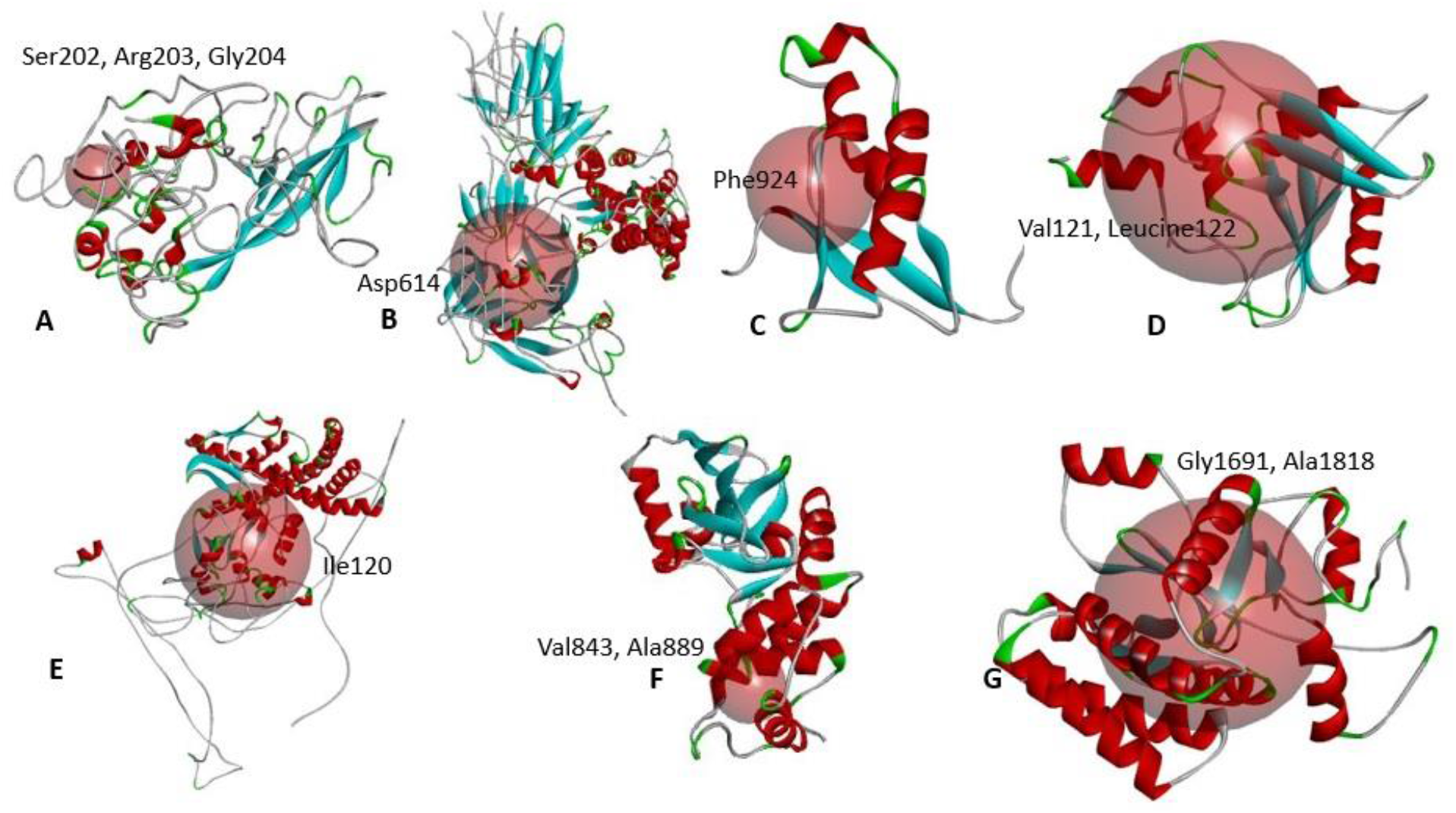
Mutation in the ligand binding site of the SARS-CoV-2 proteins.

## Discussion

COVID-19 is highly contagious and the variation in its genome could be a leading reason for this feature. Besides, to understand the origin of the strains, the exploration of the whole-genome sequencing (WGS) data of SARS-CoV-2 strains is highly necessary. [75] Insights into the mutations of SARS-CoV-2 is an important factor in developing therapeutics against the virus. [76]–[78] In this study, we investigated the variation, insertion, and deletion of the Bangladeshi SARS-CoV-2 strains. We collected the samples SARS-CoV-2/human/BGD/NIB_01/2020 and SARS-CoV-2/human/BGD/NIB-BCSIR_02/2020 from the patients who were tested as COVID-19 positive. We extracted the viral RNA from the samples and converted them to cDNA. We performed the Sanger sequencing of SARS-CoV-2/human/BGD/NIB_01/2020 and Next Generation Sequencing of SARS-CoV-2/human/BGD/NIB-BCSIR_02/2020. The total length of the genomes were 29724 and 29737 nucleotides respectively. These two genomes were submitted in both Global Initiative on Sharing All Influenza Data (GISAID) and National Center for Biotechnology Information (NCBI) databases. These two databases accepted the genomes and provided the accession number EPI_ISL_458133 from GISAID and MT509958 and MT568643 from NCBI. We investigated the possible variations of these two genome and found thirteen (13) mutations in SARS-CoV-2/human/BGD/NIB_01/2020 against the reference genome of SARS-CoV-2 (**Table 1 and Figure 2**). The mutations belonged different regions of the genome, but mostly found in the Orf11ab gene. Eight (8) mutations were found in the Orf1ab region. These mutations were 479^th^; T → A, 481^th^; C→A; 1015^th^; A → T, 2889^th^; C → T, 5098^th^; G → T; 5237^th^; C → T, 5642^th^; G → T, 8023^rd^; G → A. The mutation 93^rd^; C → T was located in the 5’ upstream region of the gene. The mutations 28733^rd^; G → A, 28734^th^; G → A and 28735^th^; G → C were located in the N gene of the genome. 23255^th^; A → G mutation was located in the S gene of the genome (**Table 1**). Among the thirteen (13) mutations, six (6) mutations, namely 93^rd^; C → T, 2889^th^; C → T, 23255^th^; A→G, 28733^rd^; G → A, 28734^th^; G → A and 28735^th^; G → C were reported in different countries according to CNCB^3^ database (**Table 1**). All the variations showed a missense effect upon structure except 2889^th^; C → T variant (**Table 1**). The other seven (7) mutations, 479^th^; T → A, 481^th^; C→A; 1015^th^; A → T, 5098^th^; G → T; 5237^th^; C → T, 5642^th^; G → T and 8023^rd^; G → A presented themselves as unique mutations in the MT509958 genome (**Table 1 and Figure 2**). Surprisingly, we didn’t find any mutation in SARS-CoV-2/human/BGD/NIB-BCSIR_02/2020. We also looked for indel profile of our assembled genomes. However, we did not find any insertion/deletion occurred into the genome. We looked for it in the rest of the genomes of SARS-CoV-2 in Bangladesh. Seven (7) deletions were found in Bangladeshi strains EPI_ISL_450343, EPI_ISL_450344, EPI_ISL_468074, EPI_ISL_514614, EPI_ISL_445213, EPI_ISL_445217 and EPI_ISL_450842. Among them, EPI_ISL_450343, EPI_ISL_450344, EPI_ISL_468074 and EPI_ISL_514614 shared common deletions. These deletions belonged to the ORF8 gene and the length ranged from 27913 to 28254 (**Table 2 and Figure 3**). We have found another deletion in the Orf7a gene whose position in the genome ranged from 27476 to 27668. This was found in an isolate from Dhaka region of Bangladesh (EPI_ISL_450842) (**Figure 3**).

Three (3) dimensional structures were built for both wild type and the mutants in order to observe the mutational impact on their corresponding proteins (**Figure 4**). We have predicted the stability of the protein structure of the corresponding variant based on energy minimization analysis. The variants 479^th^; T → A and 1015^th^; A → T were the more stable compare to the other variants. These variant’s structures consumed more energy than the wild type structure. The other variants exhibited a decrease in stability (**Table 3**). The binding site of the protein structures was analyzed to look for the location of the relevant amino acid variant. We have found that all the variants were located within the ligand binding site (**Figure 5**). Therefore, these residues could be considered very important in terms of ligand/drug binding. Ivermectin and Remdisivir were selected for the drug binding analysis. We performed molecular docking for each of the wild type and mutant structures with these two drugs and analyzed the interactions. We observed that the interacting wild type residues were replaced with different residues in after molecular docking with the drugs (**Table 4**). Here, the binding affinity was also found to be different from the wild type structure. It is to be clearly understood that only a single amino acid change from the wild type structure was responsible for these changes. From these analysis, it can be concluded that if any therapeutics are to be applied on these variants, the therapeutics might not work effectively due to the alteration of the residues in the mutant proteins.

## Conclusions

To reiterate the core of our study, we have performed whole genome sequencing of SARS-CoV-2 to identify genetic variations and then analyzed their impact on the structures of their corresponding proteins. We have also identified the insertions/deletions among all the sequenced Bangladeshi SARS-CoV-2 strains. The energy minimization and the drug binding analysis suggested that the identified mutations might have significant impact on structure and function of their target proteins. Therefore, the present study might be of great interest to the researchers/companies working to develop therapeutics against SARS-CoV-2 as well as gaining fundamental insights into pathogenesis of the virus.

## References

[1] C. Huang et al., “Clinical features of patients infected with 2019 novel coronavirus in Wuhan, China,” Lancet, vol. 395, pp. 497–506, 2020, doi: 10.1016/S0140-6736(20)30183-5.

[2] Q. Li et al., “Early Transmission Dynamics in Wuhan, China, of Novel Coronavirus–Infected Pneumonia,” N. Engl. J. Med., vol. 382, no. 13, pp. 1199–1207, Mar. 2020, doi: 10.1056/nejmoa2001316.

[3] X. Tang et al., “On the origin and continuing evolution of SARS-CoV-2,” Natl. Sci. Rev., vol. 7, no. 6, pp. 1012–1023, Jun. 2020, doi: 10.1093/nsr/nwaa036.

[4] S.-M. Chaw et al., “The origin and underlying driving forces of the SARS-CoV-2 outbreak,” J. Biomed. Sci. 2020 271, vol. 27, no. 1, pp. 1–12, Jun. 2020, doi: 10.1186/S12929-020-00665-8.

[5] J. S. Mackenzie and D. W. Smith, “COVID-19: a novel zoonotic disease caused by a coronavirus from China: what we know and what we don’t,” Microbiol. Aust., doi: 10.1071/MA20013.

[6] N. S. AlTakarli, “China’s Response to the COVID-19 Outbreak: A Model for Epidemic Preparedness and Management,” Dubai Med. J., vol. 3, no. 2, pp. 44–49, May 2020, doi: 10.1159/000508448.

[7] W. Wang et al., “Detection of SARS-CoV-2 in Different Types of Clinical Specimens,” JAMA - Journal of the American Medical Association, vol. 323, no. 18. American Medical Association, pp. 1843–1844, May 12, 2020, doi: 10.1001/jama.2020.3786.

[8] T. Burki, “The origin of SARS-CoV-2,” Lancet. Infect. Dis., vol. 20, no. 9, pp. 1018–1019, Sep. 2020, doi: 10.1016/S1473-3099(20)30641-1.

[9] L. Zou et al., “SARS-CoV-2 Viral Load in Upper Respiratory Specimens of Infected Patients,” N. Engl. J. Med., vol. 382, no. 12, pp. 1177–1179, Mar. 2020, doi: 10.1056/nejmc2001737.

[10] J. R. Larsen, M. R. Martin, J. D. Martin, P. Kuhn, and J. B. Hicks, “Modeling the Onset of Symptoms of COVID-19,” Front. Public Heal., vol. 8, p. 473, Aug. 2020, doi: 10.3389/fpubh.2020.00473.

[11] W. Guan et al., “Clinical Characteristics of Coronavirus Disease 2019 in China,” N. Engl. J. Med., vol. 382, no. 18, pp. 1708–1720, Apr. 2020, doi: 10.1056/nejmoa2002032.

[12] A. Lovato and C. de Filippis, “Clinical Presentation of COVID-19: A Systematic Review Focusing on Upper Airway Symptoms,” Ear, Nose and Throat Journal, vol. 99, no. 9. SAGE Publications Ltd, pp. 569–576, Nov. 01, 2020, doi: 10.1177/0145561320920762.

[13] C. Menni, C. H. Sudre, C. J. Steves, S. Ourselin, and T. D. Spector, “Quantifying additional COVID-19 symptoms will save lives,” The Lancet, vol. 395, no. 10241. Lancet Publishing Group, pp. e107–e108, Jun. 20, 2020, doi: 10.1016/S0140-6736(20)31281-2.

[14] N. Chen et al., “Epidemiological and clinical characteristics of 99 cases of 2019 novel coronavirus pneumonia in Wuhan, China: a descriptive study,” Lancet, vol. 395, no. 10223, pp. 507–513, Feb. 2020, doi: 10.1016/S0140-6736(20)30211-7.

[15] M. Cascella, M. Rajnik, A. Cuomo, S. C. Dulebohn, and R. Di Napoli, Features, Evaluation and Treatment Coronavirus (COVID-19). StatPearls Publishing, 2020.

[16] L. Mizrahi, H. A. Shekhidem, and S. Stern, “Age separation dramatically reduces COVID-19 mortality rate in a computational model of a large population,” Open Biol., vol. 10, no. 11, p. 200213, Nov. 2020, doi: 10.1098/rsob.200213.

[17] J. R. Goldstein and R. D. Lee, “Demographic perspectives on the mortality of COVID-19 and other epidemics,” Proc. Natl. Acad. Sci. U. S. A., vol. 117, no. 36, pp. 22035–22041, Sep. 2020, doi: 10.1073/pnas.2006392117.

[18] M. Hasanul Banna Siam, M. Mahbub Hasan, E. Raheem, M. Hasinur Rahaman Khan, M. H. Siddiqee, and M. Sorowar Hossain, “Insights into the first wave of the COVID-19 pandemic in Bangladesh: Lessons learned from a high-risk country,” 1101, doi: 10.1101/2020.08.05.20168674.

[19] C. Bonanad et al., “The Effect of Age on Mortality in Patients With COVID-19: A Meta-Analysis With 611,583 Subjects,” J. Am. Med. Dir. Assoc., vol. 21, no. 7, pp. 915–918, Jul. 2020, doi: 10.1016/j.jamda.2020.05.045.

[20] M. Brandén et al., “Residential context and COVID-19 mortality among adults aged 70 years and older in Stockholm: a population-based, observational study using individual-level data,” Lancet Heal. Longev., vol. 1, no. 2, pp. e80–e88, Nov. 2020, doi: 10.1016/s2666-7568(20)30016-7.

[21] N. D. Yanez, N. S. Weiss, J. A. Romand, and M. M. Treggiari, “COVID-19 mortality risk for older men and women,” BMC Public Health, vol. 20, no. 1, p. 1742, Dec. 2020, doi: 10.1186/s12889-020-09826-8.

[22] D. Liu et al., “Viral sepsis is a complication in patients with Novel Corona Virus Disease (COVID-19),” Med. Drug Discov., vol. 8, p. 100057, Dec. 2020, doi: 10.1016/j.medidd.2020.100057.

[23] H. C. Prescott and T. D. Girard, “Recovery from Severe COVID-19: Leveraging the Lessons of Survival from Sepsis,” JAMA - Journal of the American Medical Association, vol. 324, no. 8. American Medical Association, pp. 739–740, Aug. 25, 2020, doi: 10.1001/jama.2020.14103.

[24] A. O. Coz Yataco and S. Q. Simpson, “Coronavirus Disease 2019 Sepsis: A Nudge Toward Antibiotic Stewardship,” Chest, vol. 158, no. 5. Elsevier Inc., pp. 1833–1834, Nov. 01, 2020, doi: 10.1016/j.chest.2020.07.023.

[25] J. Beltrán-García et al., “Sepsis and Coronavirus Disease 2019: Common Features and Anti-Inflammatory Therapeutic Approaches,” Crit. Care Med., vol. 48, no. 12, pp. 1841–1844, Dec. 2020, doi: 10.1097/ccm.0000000000004625.

[26] C. Basso et al., “Pathological features of COVID-19-associated myocardial injury: a multicentre cardiovascular pathology study,” Eur. Heart J., vol. 41, no. 39, pp. 3827–3835, Oct. 2020, doi: 10.1093/eurheartj/ehaa664.

[27] K. B. Shaha, D. N. Manandhar, J. R. Cho, A. Adhikari, and M. B. K C, “COVID-19 and the heart: What we have learnt so far,” Postgraduate Medical Journal. BMJ Publishing Group, Sep. 17, 2020, doi: 10.1136/postgradmedj-2020-138284.

[28] R. D. Mitrani, N. Dabas, and J. J. Goldberger, “COVID-19 cardiac injury: Implications for long-term surveillance and outcomes in survivors,” Hear. Rhythm, vol. 17, no. 11, pp. 1984–1990, Nov. 2020, doi: 10.1016/j.hrthm.2020.06.026.

[29] C. Chen, H. Li, W. Hang, and D. W. Wang, “Cardiac injuries in coronavirus disease 2019 (COVID-19),” J. Mol. Cell. Cardiol., vol. 145, pp. 25–29, Aug. 2020, doi: 10.1016/j.yjmcc.2020.06.002.

[30] A. Tajbakhsh et al., “COVID-19 and cardiac injury: clinical manifestations, biomarkers, mechanisms, diagnosis, treatment, and follow up,” Expert Review of Anti-Infective Therapy. Taylor and Francis Ltd., 2020, doi: 10.1080/14787210.2020.1822737.

[31] B. Yang, S. Shi, M. Qin, and B. Yang, “Coronavirus Disease 2019 (COVID-19) and Cardiac Injury - Reply,” JAMA Cardiology, vol. 5, no. 10. American Medical Association, pp. 1199–1200, Oct. 01, 2020, doi: 10.1001/jamacardio.2020.2456.

[32] S. D. Unudurthi, P. Luthra, R. J. C. Bose, J. McCarthy, and M. I. Kontaridis, “Cardiac inflammation in COVID-19: Lessons from heart failure,” Life Sciences, vol. 260. Elsevier Inc., p. 118482, Nov. 01, 2020, doi: 10.1016/j.lfs.2020.118482.

[33] S. Dan, M. Pant, and S. K. Upadhyay, “The Case Fatality Rate in COVID-19 Patients With Cardiovascular Disease: Global Health Challenge and Paradigm in the Current Pandemic,” Current Pharmacology Reports, vol. 6, no. 6. Springer Science and Business Media Deutschland GmbH, pp. 315–324, Dec. 01, 2020, doi: 10.1007/s40495-020-00239-0.

[34] W. Jacobs et al., “Fatal lymphocytic cardiac damage in coronavirus disease 2019 (COVID-19): autopsy reveals a ferroptosis signature,” ESC Hear. Fail., p. ehf2.12958, Sep. 2020, doi: 10.1002/ehf2.12958.

[35] P. Aloor, “Impact of COVID-19 on chronic cardiovascular patients.”

[36] J. P. Lang, X. Wang, F. A. Moura, H. K. Siddiqi, D. A. Morrow, and E. A. Bohula, “A current review of COVID-19 for the cardiovascular specialist,” American Heart Journal, vol. 226. Mosby Inc., pp. 29–44, Aug. 01, 2020, doi: 10.1016/j.ahj.2020.04.025.

[37] D. Wang et al., “Clinical Characteristics of 138 Hospitalized Patients with 2019 Novel Coronavirus-Infected Pneumonia in Wuhan, China,” JAMA - J. Am. Med. Assoc., vol. 323, no. 11, pp. 1061–1069, Mar. 2020, doi: 10.1001/jama.2020.1585.

[38] M. L. Holshue et al., “First Case of 2019 Novel Coronavirus in the United States,” N. Engl. J. Med., vol. 382, no. 10, pp. 929–936, Mar. 2020, doi: 10.1056/nejmoa2001191.

[39] K. Prem et al., “The effect of control strategies to reduce social mixing on outcomes of the COVID-19 epidemic in Wuhan, China: a modelling study,” Lancet Public Heal., vol. 5, no. 5, pp. e261–e270, May 2020, doi: 10.1016/S2468-2667(20)30073-6.

[40] T. L. Xu et al., “China’s practice to prevent and control COVID-19 in the context of large population movement,” Infect. Dis. Poverty, vol. 9, no. 1, p. 115, Aug. 2020, doi: 10.1186/s40249-020-00716-0.

[41] M. Fugazza, “Impact of the COVID-19 pandemic on commodities exports to China,” 2020. Accessed: Dec. 10, 2020. [Online]. Available: https://www.markiteconomics.com/Public/Home/PressRelease/f69c639a88b54bc586be362511083192.

[42] M. Vinceti et al., “Lockdown timing and efficacy in controlling COVID-19 using mobile phone tracking,” EClinicalMedicine, vol. 25, p. 100457, Aug. 2020, doi: 10.1016/j.eclinm.2020.100457.

[43] J. D. Hamadani et al., “Immediate impact of stay-at-home orders to control COVID-19 transmission on socioeconomic conditions, food insecurity, mental health, and intimate partner violence in Bangladeshi women and their families: an interrupted time series,” Lancet Glob. Heal., vol. 8, no. 11, pp. e1380–e1389, Nov. 2020, doi: 10.1016/S2214-109X(20)30366-1.

[44] Z. Xu et al., “Pathological findings of COVID-19 associated with acute respiratory distress syndrome,” Lancet Respir. Med., vol. 8, no. 4, pp. 420–422, Apr. 2020, doi: 10.1016/S2213-2600(20)30076-X.

[45] L. M. Casanova, S. Jeon, W. A. Rutala, D. J. Weber, and M. D. Sobsey, “Effects of air temperature and relative humidity on coronavirus survival on surfaces,” Appl. Environ. Microbiol., vol. 76, no. 9, pp. 2712–2717, May 2010, doi: 10.1128/AEM.02291-09.

[46] S. Kumar, R. Nyodu, V. K. Maurya, and S. K. Saxena, “Morphology, Genome Organization, Replication, and Pathogenesis of Severe Acute Respiratory Syndrome Coronavirus 2 (SARS-CoV-2),” Springer, Singapore, 2020, pp. 23–31.

[47] M. Bianchi, D. Benvenuto, M. Giovanetti, S. Angeletti, M. Ciccozzi, and S. Pascarella, “Sars-CoV-2 Envelope and Membrane Proteins: Structural Differences Linked to Virus Characteristics?,” Biomed Res. Int., vol. 2020, 2020, doi: 10.1155/2020/4389089.

[48] D. Schoeman and B. C. Fielding, “Coronavirus envelope protein: current knowledge,” doi: 10.1186/s12985-019-1182-0.

[49] F. A. Rabi, M. S. Al Zoubi, G. A. Kasasbeh, D. M. Salameh, and A. D. Al-Nasser, “SARS-CoV-2 and Coronavirus Disease 2019: What We Know So Far,” Pathogens, vol. 9, no. 3, p. 231, Mar. 2020, doi: 10.3390/pathogens9030231.

[50] H. Abboud, F. Z. Abboud, H. Kharbouch, Y. Arkha, N. El Abbadi, and A. El Ouahabi, “COVID-19 and SARS-Cov-2 Infection: Pathophysiology and Clinical Effects on the Nervous System,” World Neurosurgery, vol. 140. Elsevier Inc., pp. 49–53, Aug. 01, 2020, doi: 10.1016/j.wneu.2020.05.193.

[51] S. Khan et al., “Coronaviruses disease 2019 (COVID-19): Causative agent, mental health concerns, and potential management options,” Journal of Infection and Public Health, vol. 13, no. 12. Elsevier Ltd, pp. 1840–1844, Dec. 01, 2020, doi: 10.1016/j.jiph.2020.07.010.

[52] K. A. Adedokun, A. O. Olarinmoye, A. O. Olarinmoye, J. O. Mustapha, and R. T. Kamorudeen, “A close look at the biology of SARS-CoV-2, and the potential influence of weather conditions and seasons on COVID-19 case spread,” Infect. Dis. Poverty, vol. 9, no. 1, p. 77, Jun. 2020, doi: 10.1186/s40249-020-00688-1.

[53] S. Ludwig and A. Zarbock, “Coronaviruses and SARS-CoV-2: A Brief Overview,” Anesth. Analg., pp. 93–96, 2020, doi: 10.1213/ANE.0000000000004845.

[54] M. Cevik, M. Tate, O. Lloyd, A. E. Maraolo, J. Schafers, and A. Ho, “SARS-CoV-2, SARS-CoV, and MERS-CoV viral load dynamics, duration of viral shedding, and infectiousness: a systematic review and meta-analysis,” The Lancet Microbe, vol. 0, no. 0, Nov. 2020, doi: 10.1016/s2666-5247(20)30172-5.

[55] D. Wu, T. Wu, Q. Liu, and Z. Yang, “The SARS-CoV-2 outbreak: What we know,” Int. J. Infect. Dis., vol. 94, pp. 44–48, 2020, doi: 10.1016/j.ijid.2020.03.004.

[56] Z. Abdelrahman, M. Li, and X. Wang, “Comparative Review of SARS-CoV-2, SARS-CoV, MERS-CoV, and Influenza A Respiratory Viruses,” Frontiers in Immunology, vol. 11. Frontiers Media S.A., p. 552909, Sep. 11, 2020, doi: 10.3389/fimmu.2020.552909.

[57] M. Fani, A. Teimoori, and S. Ghafari, “Comparison of the COVID-2019 (SARS-CoV-2) pathogenesis with SARS-CoV and MERS-CoV infections,” Future Virology, vol. 15, no. 5. Future Medicine Ltd., pp. 317–323, May 01, 2020, doi: 10.2217/fvl-2020-0050.

[58] Z. Zhu, X. Lian, X. Su, W. Wu, G. A. Marraro, and Y. Zeng, “From SARS and MERS to COVID-19: A brief summary and comparison of severe acute respiratory infections caused by three highly pathogenic human coronaviruses,” Respiratory Research, vol. 21. 1. BioMed Central Ltd, p. 224, Aug. 27, 2020, doi: 10.1186/s12931-020-01479-w.

[59] Y. Chen, Q. Liu, and D. Guo, “Emerging coronaviruses: Genome structure, replication, and pathogenesis,” Journal of Medical Virology, vol. 92, no. 4. John Wiley and Sons Inc., pp. 418–423, Apr. 01, 2020, doi: 10.1002/jmv.25681.

[60] A. Wu et al., “Genome Composition and Divergence of the Novel Coronavirus (2019-nCoV) Originating in China,” Cell Host Microbe, vol. 27, no. 3, pp. 325–328, Mar. 2020, doi: 10.1016/j.chom.2020.02.001.

[61] A. R. Fehr and S. Perlman, “Coronaviruses: An overview of their replication and pathogenesis,” in Coronaviruses: Methods and Protocols, vol. 1282, Springer New York, 2015, pp. 1–23.

[62] J. Hellewell et al., “Feasibility of controlling COVID-19 outbreaks by isolation of cases and contacts,” Lancet Glob. Heal., vol. 8, no. 4, pp. e488–e496, Apr. 2020, doi: 10.1016/S2214-109X(20)30074-7.

[63] R. Lu et al., “Genomic characterisation and epidemiology of 2019 novel coronavirus: implications for virus origins and receptor binding,” Lancet, vol. 395, no. 10224, pp. 565–574, Feb. 2020, doi: 10.1016/S0140-6736(20)30251-8.

[64] M. Moniruzzaman et al., “Coding-Complete Genome Sequence of SARS-CoV-2 Isolate from Bangladesh by Sanger Sequencing,” Microbiol. Resour. Announc., vol. 9, no. 28, Jul. 2020, doi: 10.1128/mra.00626-20.

[65] Heracle BioSoft, “DNA Sequence Assembler v4 (2013).” https://www.dnabaser.com/ (accessed Dec. 10, 2020).

[66] T. G. Burland, “DNASTAR’s Lasergene sequence analysis software.,” Methods Mol. Biol., vol. 132, pp. 71–91, 2000, doi: 10.1385/1-59259-192-2:71.

[67] J. L. A. Paijmans et al., “Sequencing single-stranded libraries on the Illumina NextSeq 500 platform.” Accessed: Dec. 10, 2020. [Online]. Available: https://www.illumina.com/systems/sequencing-.

[68] M. Johnson, I. Zaretskaya, Y. Raytselis, Y. Merezhuk, S. McGinnis, and T. L. Madden, “NCBI BLAST: a better web interface.,” Nucleic Acids Res., vol. 36, no. Web Server issue, pp. 5–9, Jul. 2008, doi: 10.1093/nar/gkn201.

[69] Y. Shu and J. McCauley, “GISAID: Global initiative on sharing all influenza data – from vision to reality,” Eurosurveillance, vol. 22, no. 13. European Centre for Disease Prevention and Control (ECDC), p. 1, Mar. 30, 2017, doi: 10.2807/1560-7917.ES.2017.22.13.30494.

[70] T. J. Carver, K. M. Rutherford, M. Berriman, M.-A. Rajandream, B. G. Barrell, and J. Parkhill, “ACT: the Artemis comparison tool,” Bioinformatics, vol. 21, no. 16, pp. 3422–3423, Aug. 2005, doi: 10.1093/bioinformatics/bti553.

[71] D. E. Kim, D. Chivian, and D. Baker, “Protein structure prediction and analysis using the Robetta server,” Nucleic Acids Res., vol. 32, no. WEB SERVER ISS., pp. W526–W531, Jul. 2004, doi: 10.1093/nar/gkh468.

[72] E. Lindahl, B. Hess, and D. van der Spoel, “GROMACS 3.0: A package for molecular simulation and trajectory analysis,” Journal of Molecular Modeling, vol. 7, no. 8. Springer, pp. 306–317, 2001, doi: 10.1007/S008940100045.

[73] O. Trott and A. J. Olson, “AutoDock Vina: Improving the speed and accuracy of docking with a new scoring function, efficient optimization, and multithreading,” J. Comput. Chem., vol. 31, no. 2, p. NA-NA, 2009, doi: 10.1002/jcc.21334.

[74] T. G. Bell and A. Kramvis, “Fragment merger: An online tool to merge overlapping long sequence fragments,” Viruses, vol. 5, no. 3, pp. 824–833, 2013, doi: 10.3390/v5030824.

[75] I. Ahammad et al., “Comparative Genomic Study for Revealing the Complete Scenario of COVID-19 Pandemic in Bangladesh,” medRxiv, p. 2020.11.27.20240002, Nov. 2020, doi: 10.1101/2020.11.27.20240002.

[76] B. Dearlove et al., “A SARS-CoV-2 vaccine candidate would likely match all currently circulating variants,” Proc. Natl. Acad. Sci. U. S. A., vol. 117, no. 38, pp. 23652–23662, Sep. 2020, doi: 10.1073/pnas.2008281117.

[77] I. Ahammad and S. S. Lira, “Designing a novel mRNA vaccine against SARS-CoV-2: An immunoinformatics approach,” Int. J. Biol. Macromol., vol. 162, pp. 820–837, Nov. 2020, doi: 10.1016/j.ijbiomac.2020.06.213.

[78] M. I. Abdelmageed et al., “Design of a Multiepitope-Based Peptide Vaccine against the e Protein of Human COVID-19: An Immunoinformatics Approach,” Biomed Res. Int., vol. 2020, 2020, doi: 10.1155/2020/2683286.

